# VAMPr: <u>VA</u>riant <u>M</u>apping and <u>P</u>rediction of antibiotic <u>r</u>esistance via explainable features and machine learning

**DOI:** 10.1101/537381

**Authors:** Jiwoong Kim, David E Greenberg, Reed Pifer, Shuang Jiang, Guanghua Xiao, Samuel A Shelburne, Andrew Koh, Yang Xie, Xiaowei Zhan

**Affiliations:** Quantitative Biomedical Research Center, Department of Clinical Sciences, University of Texas Southwestern Medical Center, Dallas, Texas, 75390, USA; Department of Internal Medicine, University of Texas Southwestern Medical Center, Dallas, Texas, 75390, USA; Department of Microbiology, University of Texas Southwestern Medical Center, Dallas, Texas, 75390, USA; Department of Statistical Science, Southern Methodist University, Dallas, TX 75275, USA; Harold C. Simmons Cancer Center, University of Texas Southwestern Medical Center, Dallas, Texas, 75390, USA; Department of Bioinformatics, University of Texas Southwestern Medical Center, Dallas, Texas, 75390, USA; Department of Infectious Diseases and Genomic Medicine, University of Texas MD Anderson Cancer Center, Houston, Texas, USA; Department of Pediatrics, University of Texas Southwestern Medical Center, Dallas, Texas, 75390, USA; Center for Genetics of Host Defense, University of Texas Southwestern Medical Center, Dallas, Texas, 75390, USA

## Abstract

Antimicrobial resistance (AMR) is an increasing threat to public health. Current methods of determining AMR rely on inefficient phenotypic approaches, and there remains incomplete understanding of AMR mechanisms for many pathogen-antimicrobial combinations. Given the rapid, ongoing increase in availability of high density genomic data for a diverse array of bacteria, development of algorithms that could utilize genomic information to predict phenotype could both be useful clinically and assist with discovery of heretofore unrecognized AMR pathways. To facilitate understanding of the connections between DNA variation and phenotypic AMR, we developed a new bioinformatics tool, variant mapping and prediction of antibiotic resistance (VAMPr), to (1) derive gene ortholog-based sequence features for variants; (2) interrogate these explainable gene-level variants for their known or novel associations with AMR; and (3) build accurate models to predict AMR based on whole genome sequencing data. Following the Clinical & Laboratory Standards Institute (CLSI) guidelines, we curated the publicly available sequencing data for 3,393 bacterial isolates from 9 species along with AMR phenotypes for 29 antibiotics. We detected 14,615 variant genotypes and built 93 association and prediction models. The association models confirmed known genetic antibiotic resistance mechanisms, such as blaKPC and carbapenem resistance consistent with the accurate nature of our approach. The prediction models achieved high accuracies (mean accuracy of 91.1% for all antibiotic-pathogen combinations) internally through nested cross validation and were also validated using external clinical datasets. The VAMPr variant detection method, association and prediction models will be valuable tools for AMR research for basic scientists with potential for clinical applicability.

## INTRODUCTION

Antimicrobial resistance (AMR) is an urgent worldwide threat (1). Decreased efficacy of antibiotics can lead to prolonged hospitalization and increased mortality (2). Current phenotypic methods for determining whether an isolate is sensitive or resistant to a particular antibiotic can, in some instances, take days resulting in delays in providing effective therapy (3). Targeted methods for AMR determination, such as PCR, are limited in that they identify only a subset of resistant genes and therefore do not provide a full explanation for a particular resistance phenotype (4).

Next-generation sequencing (NGS) technology enabling whole genome sequencing (WGS) of bacterial isolates is now both inexpensive and widely-used (5). We have previously shown that NGS can identify AMR determinants for a limited number of β-lactam antimicrobials and that genotype correlated well with classic phenotypic testing (6). However, that study focused on a narrow set of both antibiotics and pathogens because the links between genotype and phenotype are relatively well understood for those antibiotic/pathogen combinations. Other groups have utilized NGS data to identify the presence of genes or short nucleotide sequences that confer resistance in a variety of pathogens (7-9). Mechanisms of AMR for many pathogen-antibiotic combinations are not well delineated which hinders development of genotypic-phenotypic associations. In order to more fully explore genotypic prediction of antibiotic resistance and build upon our previous efforts, we have developed novel methods for utilizing NGS data to better 1) characterize amino-acid based variant features, 2) expand the knowledge base of genetic associations with AMR, and 3) construct accurate prediction models for determining phenotypic resistance from NGS data in a broad array of pathogen-antibiotic combinations. This algorithm, called VAMPr (**VA**riant **M**apping and **P**rediction of antibiotic **r**esistance), was built utilizing a large dataset of bacterial genomes from the NCBI SRA along with paired antibiotic susceptibility data from the NCBI BioSample Antibiogram. VAMPr utilizes two different approaches, association models and prediction models, to assess genotype-phenotype relationship. In the association analysis, data-driven association models utilizing a gene ortholog approach were constructed. This allowed for unbiased screening of genotype and phenotype across a broad array of bacterial isolates. In the prediction analysis, we utilized a machine learning algorithm to develop prediction models that take NGS data and predict resistance for every pathogen-drug combination. These approaches not only confirmed known genetic mechanisms of antibacterial resistance, but also identified potentially novel or underreported correlates of resistance.

## MATERIAL AND METHODS

### Overview of VAMPr

The VAMPr workflow is depicted in Figure 1. Publicly available bacterial genomes from the NCBI Short Read Archive (SRA) and paired antibiotic susceptibility data from the NCBI BioSample Antibiograms project were downloaded. In order to identify bacterial genetic variants, *de novo* assembly was performed and assembled scaffolds were aligned to the Antimicrobial Resistance (AMR) KEGG orthology database (KO)(10). Through this process KO-based sequence variants were identified. CLSI breakpoints were used to determine the antibiotic phenotype (sensitive versus resistant; isolates with intermediate susceptibility were not included for analysis)(11). Finally, factoring both genetic variants and antibiotic resistance phenotypes, association and prediction models were constructed. These models are available to the research community through our website (see **Availability**).

**Figure 1.**
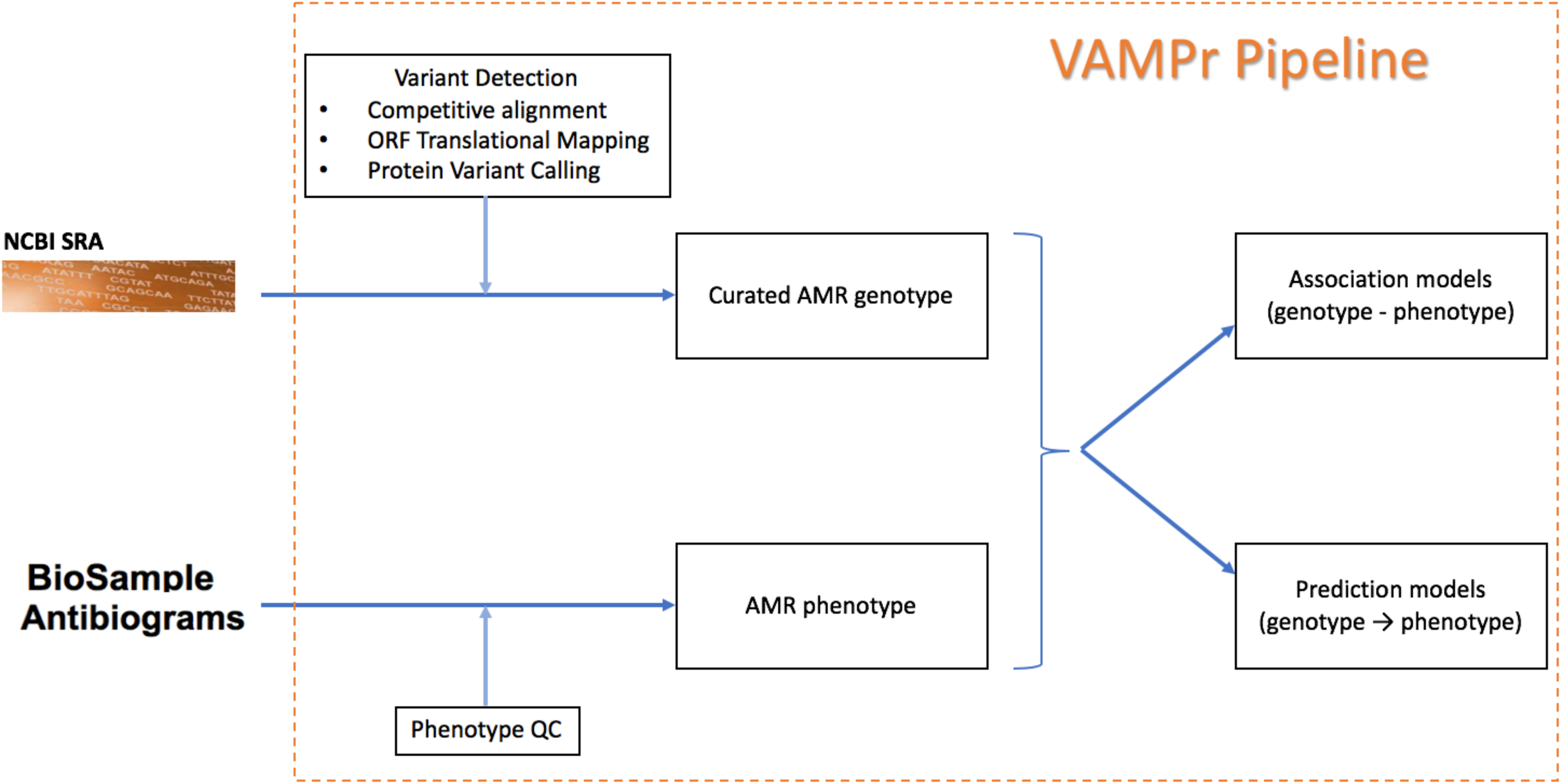
Overview of the VAMPr workflow. The VAMPr pipeline processed sequence data from the NCBI Short Read Achieve (SRA) and NCBI BioSample Antibiograms for phenotypes. The curated AMR genotypes and AMR phenotypes were used to create both association and prediction models.

### Data Acquisition and Creation of AMR Protein Database

Bacterial isolates with antibiotic susceptibility data were identified in the NCBI BioSample Antibiograms database. Isolates were identified by querying “antibiogram[filter]” in the National Center for Biotechnology Information (NCBI) (NCBI Resource Coordinators, 2018) BioSample. The linked sequencing data was downloaded from the NCBI Sequence Read Archive (SRA). Finally, the antibiogram tables in the NCBI BioSample were downloaded using NCBI API. Minimum inhibitory concentration (MIC) values and reported antibiotic susceptibility data were recorded and checked for accuracy according to CLSI guidelines (11). MIC values that were clearly mis-annotated were removed. For the purposes of this analysis, isolates that were intermediate for any particular drug were excluded. In addition, any bacterial isolate reported as both resistant and susceptible was excluded from analysis.

A reference database consisting of KO genes with gene-based variants was created that included both AMR protein sequences as well as AMR-like protein sequences (decoy sequences). The AMR-like sequences are from genes known to not be involved in antibiotic resistance and have been shown to improve variant calling accuracies (12). To create the AMR protein database, a list of Kyoto Encyclopedia of Genes and Genomes (KEGG) orthology (KO) involved in antimicrobial resistance (AMR) (**Supplementary Figure 1**) was created. The protein sequences linked to AMR KOs by KEGG API and UniProt ID mapping (13) were downloaded from the UniProt database. These sequences were designated as AMR protein sequences. Further, protein sequences from KOs not related to AMR were also aligned to AMR protein sequences. AMR-like protein sequences were defined as those protein sequences with 80% identical amino acid alignment. The union of AMR protein sequences and AMR-like protein sequences formed the AMR protein database which was utilized in all comparative alignment steps.

To facilitate the identification of variants, AMR protein sequences were clustered based on sequence identities using CD-HIT(14). For each cluster, multiple sequence alignment (MSA) steps were used to determine cluster consensus sequences (CCS) using MAFFT (15). Finally, bacterial isolate protein sequences were compared to CCSs to identify the variants (see next section and **Supplementary: Derive explainable KO gene-based sequence variants**).

### Characterization of AMR Variants

We developed an algorithm to characterize the AMR-related variants at the protein level (**Supplementary materials**). For each individual bacterial isolate, *de novo* genome assembly was performed using SPAdes (16). Open reading frame (ORF)s were identified, converted to amino acid sequences, and, protein BLAST of the sequences using the aforementioned AMR protein database was performed. The same query sequence was aligned to both AMR and AMR-like reference protein sequences using Diamond (17). After comparative alignments and removal of less than 80% identical amino acids, only alignments best matched to the AMR reference sequences were included (see **Supplementary: Comparative alignments** for filters on E-values, bit-scores and fraction of identical amino acids and **Supplementary Figure 2**). Finally, the aligned protein scaffold sequences were compared to the CCS to define a “normal” protein versus a variant. For example, given a perfect match, an isolate is designated as carrying the KO gene and thus denoted as normal. In contrast, if there were mismatched amino acids within a CCS alignment, these would be deemed as novel variants, and in such cases, the detected variants would have the following nomenclature: KO number, KO cluster number, sequence variant types and their details (substitution, insertion, deletion). More details are provided in **Supplementary Figure 3** and **Supplementary materials**.

### VAMPr association model to characterize variants

To quantitatively assess the association between KO-based sequence variants and antibiotic resistance phenotypes, an association model for each species-antibiotic combination was created. In total, 52,479 associations between variants and antibiotic resistance were evaluated. Specifically, a 2-by-2 contingency table for all isolates based on carrier/non-carrier status of the variant and susceptible/resistant phenotypes was generated and the odds ratio and p-values based on Fisher’s exact test were calculated in R 3.4.4 (18) and adjusted for false discovery rate based on Benjamin-Hochberg procedure (19). The fraction of resistant strains stratified by the variants’ carrying status was visualized in bar plots.

### VAMPr prediction model for antibiotic resistance

Prediction models for each species-antibiotic combination were developed. KO-based sequence variants were designated as features and curated antibiotic resistant phenotypes as labels. For each species-antibiotics combination, an optimal prediction model with tuned hyperparameters was generated. A gradient boosting tree approach was utilized, given its accurate performance profile and efficient implementation (20). Nested cross-validation (CV) was used to report unbiased prediction performance (21,22). The outer CV was 10-fold and the averaged prediction metrics including accuracy are reported (Table 1); the inner CV was 5-fold and all inner folds were used for hyper-parameter tuning based on prediction accuracy. The default search space hyperparameters were chosen as follows: the number of rounds (the number of trees) was 50, 100, 500 or 1000; the maximum allowed depth of trees was 16 or 64; the learning rate was from 0.025 or 0.05; the minimum loss reduction required to allow further partition of the trees was 0; the fraction of features used for constructing each tree was 0.8; the fraction of isolates used for constructing each tree was 0.9; and the minimum weight for each child tree was 0. The reported performance metrics included accuracy, F1-score, and area under the receiver operating characteristic curve (AUROC) (21). We assessed the prediction accuracy using an independent dataset of bacterial isolates recovered from cancer patients with bloodstream infections (6). In this study (11), all isolates were genetically (whole-genome sequence) and phenotypically (antibiotic susceptibility testing by broth microdilution assays) profiled. We followed the same aforementioned genotype and phenotype processing steps. The detected KO-based variants were used as predictors and the lab-measured antibiotic resistance phenotypes were used as the gold standard. Performance metrics were calculated as described above.

**Table 1.** Prediction metrics for 32 VAMPr prediction models. Among 93 prediction models, we listed the top 32 models that have the mean prediction accuracies higher than 95%. The isolate and variant counts derived from sequencing were used to build the prediction model using gradient boosting tree algorithms. The accuracy is reported using nested cross validation approach. The 10-fold outer cross validation were used to report accuracy and the 5-fold inner cross validation was used for hyperparameter tuning.

| Species | Antibiotics | Isolate counts | Variant Counts | Fraction of Resistant Isolates | Accuracy |
| --- | --- | --- | --- | --- | --- |
| *Streptococcus pneumoniae* | tetracycline | 315 | 1,321 | 6.0% | 100.0% |
| *Streptococcus pneumoniae* | meropenem | 173 | 1,218 | 5.8% | 100.0% |
| *Streptococcus pneumoniae* | amoxicillin | 171 | 1,208 | 2.3% | 100.0% |
| *Escherichia coli* | kanamycin | 75 | 827 | 13.3% | 100.0% |
| *Staphylococcus aureus* | tetracycline | 31 | 540 | 9.7% | 100.0% |
| *Staphylococcus aureus* | clindamycin | 24 | 479 | 37.5% | 100.0% |
| *Salmonella enterica* | trimethoprim-sulfamethoxazole | 1,349 | 1,620 | 0.7% | 99.6% |
| *Streptococcus pneumoniae* | cefuroxime | 178 | 1,221 | 9.6% | 99.4% |
| *Salmonella enterica* | cefoxitin | 1,291 | 1,483 | 15.9% | 99.3% |
| *Salmonella enterica* | chloramphenicol | 1,337 | 1,510 | 3.3% | 99.3% |
| *Salmonella enterica* | amoxicillin-clavulanic acid | 1,285 | 1,476 | 19.8% | 99.1% |
| *Salmonella enterica* | kanamycin | 1,036 | 1,373 | 9.4% | 98.9% |
| *Salmonella enterica* | ceftiofur | 1,340 | 1,489 | 19.0% | 98.8% |
| *Salmonella enterica* | ceftriaxone | 1,345 | 1,536 | 19.3% | 98.7% |
| *Salmonella enterica* | tetracycline | 1,339 | 1,518 | 53.0% | 98.6% |
| *Salmonella enterica* | ampicillin | 1,349 | 1,620 | 33.2% | 98.5% |
| *Streptococcus pneumoniae* | clindamycin | 316 | 1,323 | 3.5% | 98.4% |
| *Klebsiella aerogenes* | tobramycin | 58 | 1,014 | 5.2% | 98.3% |
| *Salmonella enterica* | gentamicin | 1,333 | 1,507 | 12.3% | 98.0% |
| *Klebsiella pneumoniae* | cefotaxime | 294 | 1,808 | 97.3% | 97.6% |
| *Acinetobacter baumannii* | doripenem | 204 | 2,027 | 25.0% | 97.6% |
| *Streptococcus pneumoniae* | trimethoprim-sulfamethoxazole | 275 | 1,143 | 5.8% | 96.4% |
| *Acinetobacter baumannii* | amikacin | 465 | 3,427 | 9.7% | 96.4% |
| *Acinetobacter baumannii* | tetracycline | 261 | 3,096 | 23.0% | 96.2% |
| *Escherichia coli* | amikacin | 269 | 1,750 | 6.3% | 95.9% |
| *Enterobacter cloacae* | doripenem | 44 | 1,167 | 47.7% | 95.8% |
| *Escherichia coli* | imipenem | 143 | 1,049 | 22.4% | 95.8% |
| *Acinetobacter baumannii* | ciprofloxacin | 367 | 3,257 | 73.3% | 95.4% |
| *Streptococcus pneumoniae* | erythromycin | 317 | 1,323 | 28.1% | 95.3% |
| *Enterobacter cloacae* | tobramycin | 66 | 1,468 | 39.4% | 95.2% |
| *Escherichia coli* | ampicillin | 348 | 2,180 | 92.0% | 95.1% |
| *Klebsiella pneumoniae* | ertapenem | 318 | 1,983 | 86.2% | 95.0% |

## RESULTS

### Construction of NCBI datasets of curated genotypes and phenotypes

Focusing on the isolates reported in the NCBI Antibiogram database, we retrieved 4,515 bacterial whole genome sequence datasets (Illumina platform). Sequence reads were *de novo* assembled and aligned to Multi Locus Sequence Typing (MLST) databases to validate reported bacteria species identification(23). 1,100 isolates were excluded from analysis because of inaccurate species identification. Our final analysis cohort included 3,393 isolates representing 9 species: *Salmonella enterica* (1349 isolates), *Acinetobacter baumannii* (772), *Escherichia coli* (350), *Klebsiella pneumoniae* (344), *Streptococcus pneumoniae* (317), *Pseudomonas aeruginosa* (83), *Enterobacter cloacae* (79), *Klebsiella aerogenes* (68), and *Staphylococcus aureus* (31). A total of 38,871 MIC values were reported for 29 different antibiotics. (**Supplementary Table 1**) In total, there were 38,248 individual pathogen-drug data points identified (Figure 2A).

**Figure 2.**
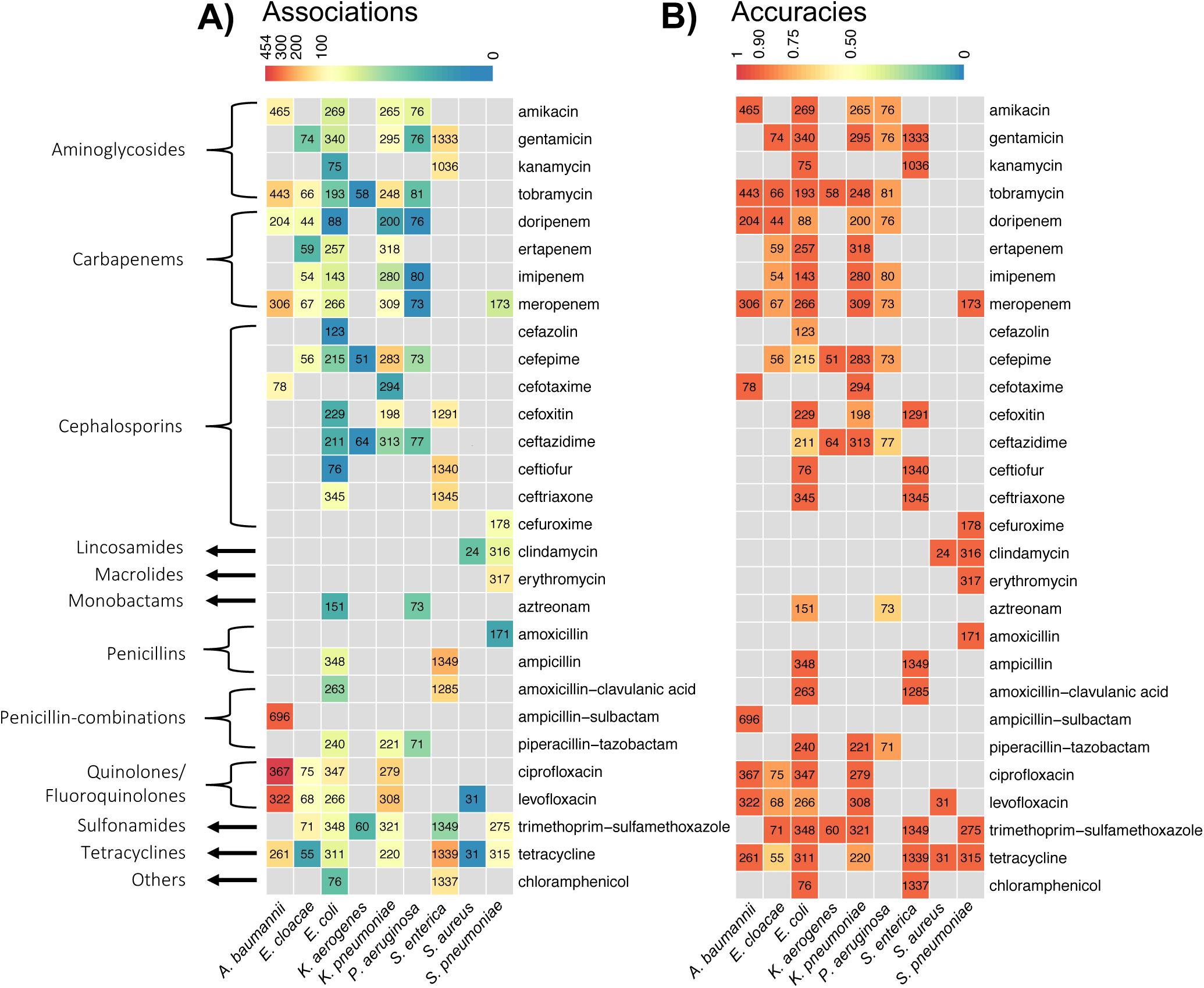
Summary of significant variant associations and prediction accuracies from 93 species-antibiotics combinations. Both heatmaps display the counts of curated isolates by the combination of 9 bacteria species and 29 antibiotics from 13 drug categories. The boxes without a number indicates that no isolates were available for this particular bacteria species and antibiotic combination. A) the color of the boxes indicates the number of gene-antibiotic resistance associations with nominal p-values <0.05 from VAMPr association models; B) the color indicates cross-validated prediction accuracies from VAMPr prediction models.

After curation, we analyzed isolates with *de novo* assembled genome and MIC values, and this dataset included 93 species/antibiotic combinations for building association and prediction models (detailed in next 3 sections). The fraction of resistant isolates for any given bacteria and antibiotic varied greatly (the median fraction of resistant isolates was 50.0%). For example, for *Salmonella enterica* and trimethoprim-sulfamethoxazole, the fraction of resistant isolates was 0.6% while for *Klebsiella pneumoniae* and cefazolin, the fraction of resistant isolates was 97.3%. This dataset was used in both the association and prediction models.

### Characterization of explainable AMR sequence variants

We curated a list of 537 Antimicrobial Resistance (AMR) KEGG ortholog (KO) genes (**Supplementary Table 2**) and then identified the corresponding UniRef protein sequences (a total of 298,760 sequences). Protein sequences were then clustered (using a minimal sequence similarity of 0.7). This resulted in 96,462 KO gene clusters to serve as a reference AMR protein sequence database. Next, we analyzed 3,393 *de novo* assembled genomes, identified the gene locations on the assembled genomes, and aligned the gene sequences to the reference AMR protein sequence database. Based on the alignment results and stringent filtering, we can identify AMR genes for each isolate. Finally, the AMR genes were examined for the presence of mutations (e.g. amino acid substitutions) using multiple sequence alignment software. We nominated an identifier format to represent the sequences. For example, K01990.129|290|TNIID indicates that the 129^th^ cluster of K01990 KO gene has mutation starting from its 290^th^ amino acid from threonine (T) and asparagine (N) to isoleucine (I) and aspartic acid (D).

### Association models between sequence variants and antibiotic resistance phenotypes retain accuracy

We interrogated the strength of the association model between genetic variants and antibiotic susceptibility phenotypes for each bacterial species and antibiotic combination. For a number of pathogen-antibiotic pairs, the association model accuracy was greater than 95% (ranged from 69.6% for *Pseudomonas aeruginosa*-aztreonam to 100.0% for *Streptococcus pneumoniae*-tetracycline; mean accuracy was 91.1%) (Figure 2B). Utilizing contingency tables of variant carrying status and resistance phenotypes with the appropriate statistical analysis (odds ratios and p-values from Fisher’s exact tests), we examined a subset of 5,359 associations with false discovery rates less than 0.05. In many instances, a significantly strong association confirmed an expected antibiotic resistance mechanism (Figure 3). For example, the sequence variant K18768.0 represents β-lactamase (Bla) encoding gene *bla*_*KPC*_, the *K. pneumoniae* carbapenemase whose presence is significantly associated with resistance to meropenem in *K. pneumoniae* (P-value <0.0001) (9)(Fig. 3A). Variant K18093.13 is *oprD*, a major porin responsible for uptake of carbapenems in *Pseudomonas* (24). Loss of porin activity by *Pseudomonas* is well known to result in carbapenem resistance (25) in this pathogen, and absence of wild-type *oprD* is strongly associated with imipenem resistance (P-value <0.0001) (Fig. 3B). Other examples (*OXA-1* and *aac-(6’)-lb*) of strong associations are illustrated in Fig. 3C and 3D.

**Figure 3.**
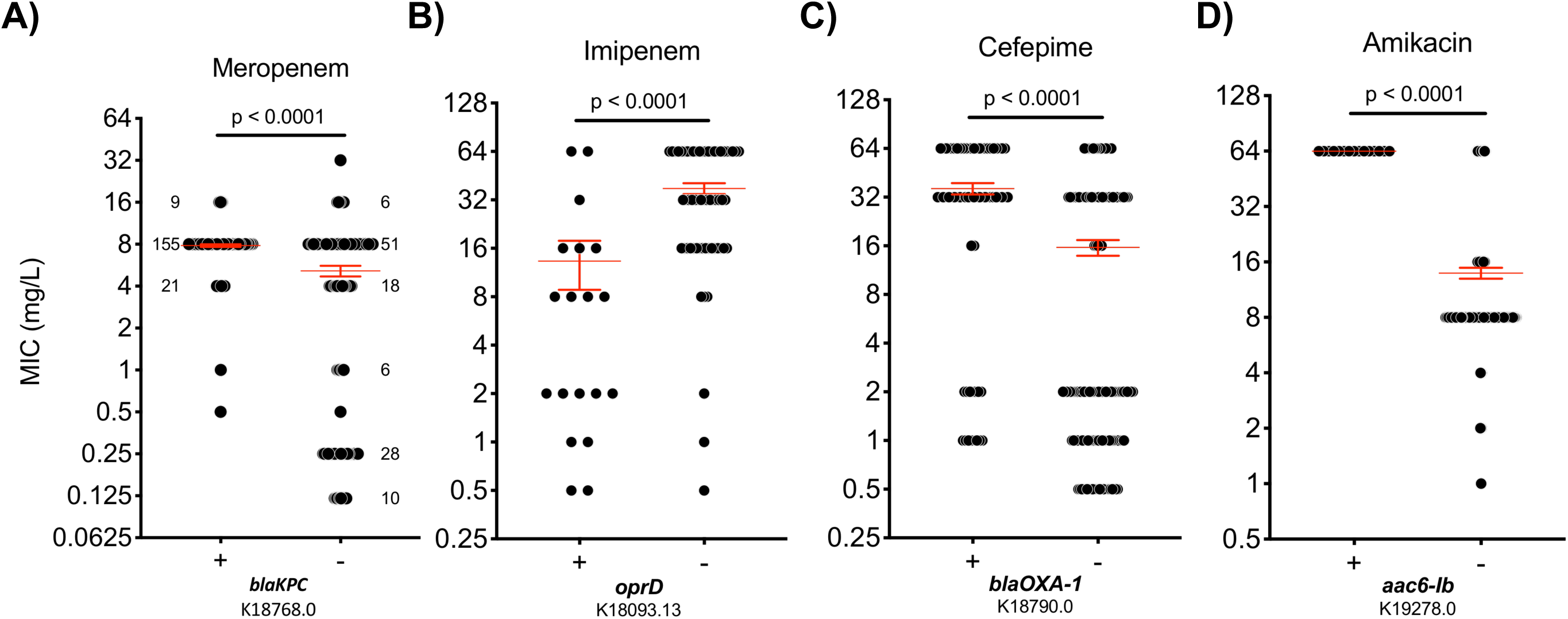
Examples of variant-phenotype relationships determined by the association models. (A) K18768.0 indicates *blaKPC*, the *K. pneumoniae* carbapenemase. The presence of blaKPC is associated with resistance to ceftazidime in *K. pneumoniae* as shown. The numbers in the plots represent the frequency of certain MIC values. Numbers in the plot represent total number of isolates with the given MIC value. (B) K18093.13 is *oprD*, an imipenem/basic amino acid-specific outer membrane pore; absence of *oprD* is associated with resistance to imipenem in *P. aeruginosa.* (C) K18790.0 represents *blaOXA-1*, the beta-lactamase class D OXA-1. Its presence is associated with resistance to cefepime in *E. coli*. (D) K19278.0 is *aac6-lb* gene. The presence of this variant is associated with amikacin resistance in *A. baumannii*. The “+” and “-” sign in the X-axis represent whether the wild-type gene exists or not. The red horizontal lines mark the mean and standard error of the groupwise MIC measurements. Each gray dot represents an MIC value. P-values are calculated based on Fisher’s exact test.

### Antibiotic Resistance Prediction Models Developed Utilizing Machine Learning

Our association studies demonstrated the accuracy of our genotypic approach for known AMR elements. To begin to explore the capacity of our approach to take sequence data and generate robust prediction, we first developed 93 different prediction models using the VAMPr pipeline. The most promising prediction models were based on an extreme boosting tree algorithm and all hyper-parameters were fine-tuned in the inner 5-fold cross validation. Other prediction models (e.g. elastic net (26), support vector machines (27), 3-layer neural network (28), and adaptive boosting (29)) were evaluated but did not exhibit superior prediction performances (**Supplementary Figure 4**). For all models, we used nested cross validation to report prediction performance metrics (Table 1). Among 93 models, half had prediction accuracies greater than 90%. The pathogen-antibiotic combinations that displayed the highest accuracy were for *Streptococcus pneumoniae* and clindamycin (100.0%), meropenem (100.0%), and tetracycline (100.0%), and *Escherichia coli* and kanamycin (100.0%). 11 prediction models for *Salmonella enterica* and our accuracies tended to be higher for this organism (minimal prediction accuracy is 98.0%) likely due to the larger dataset of *Salmonella enterica* isolates. A similar trend was also suggested by observing the performance of the models in *Acinetobacter baumannii*.

### Validation of the VAMPr prediction model using an external dataset

To validate the prediction performance of VAMPr, we utilized 13 *Enterobacter cloacae*, 31 *Escherichia coli*, 24 *Klebsiella pneumoniae* and 21 *Pseudomonas aeruginosa* isolates that were genetically and phenotypically profiled in a prior study but not present in the NCBI Antibiogram database (6). All isolates had been previously tested against 3 antibiotics (cefepime, ceftazidime, and meropenem). Importantly, approximately 62%, 15%, 28% and 31% of the discovered variants of these strains, respectively, were not detected in the NCBI isolates. As these variants are specific to the validation datasets, their roles in antibiotic resistance could not be modelled by the NCBI datasets. In Figure 4, we show three prediction results with the highest AUROC (area under the receiver operator characteristics) values, as well as the important genetic variants that frequently appear in the gradient boosting tree models. In the *Escherichia coli* and meropenem model, VAMPr reached 1.0 AUROC (Figure 4A) and the most important predictor was the presence of the *blaNDM* gene (New Dehli metallo-beta-lactamase; Class B). VAMPr had a similarly high prediction performance for *Klebsiella pneumoniae* and ceftazidime (Figure 4B). This model also has an AUROC value of 0.99 and the significant predictors were the presence of KPC (*Klebsiella pneumoniae* carbapenemase) and the presence of wildtype ddl; D-alanine-D-alanine ligase (in 4 isolates, variants of *ddl* were associated with sensitivity to ceftazidime). In Figure 4C, the prediction model for *Pseudomonas aeruginosa* and meropenem is 0.95, and three significant predictors were *ebr* (small multidrug resistance pump), *mexA* (membrane fusion protein, multidrug efflux system) and *oprD* (imipenem/basic amino acid-specific outer membrane pore). Among all bacteria and antibiotic combinations, the minimal AUROC values for all VAMPr prediction models is 0.70 (Table 2). These results suggest that the VAMPr prediction models identify both known AMR-related genes as well as genes or variants that are not currently considered as contributing to resistance.

**Table 2.** External validation of VAMPr prediction model. The external dataset includes 31 *Escherichia coli*, 24 *Klebsiella pneumoniae* and 21 *Pseudomonas aeruginosa* isolates. All isolates were tested against 3 antibiotics (cefepime, ceftazidime and meropenem). We reported the accuracy as the fraction of correct predictions, and the AUC (area under the curve) represents the area under the operator-receiver characteristic. The AUC value is n/a for E. cloacae as all 13 isolates are susceptible to meropenem.

| Species | Antibiotics | Isolate counts | Accuracy | AUC |
| --- | --- | --- | --- | --- |
| *Enterobacter cloacae* | cefepime | 11 | 100.0% | 1.00 |
| *Enterobacter cloacae* | meropenem | 13 | 92.3% | n/a |
| *Escherichia coli* | cefepime | 30 | 63.3% | 0.70 |
| *Escherichia coli* | ceftazidime | 28 | 78.6% | 0.88 |
| *Escherichia coli* | meropenem | 31 | 96.8% | 1.00 |
| *Klebsiella pneumoniae* | cefepime | 24 | 70.8% | 0.87 |
| *Klebsiella pneumoniae* | ceftazidime | 24 | 66.7% | 0.99 |
| *Klebsiella pneumoniae* | meropenem | 23 | 78.3% | 1.00 |
| *Pseudomonas aeruginosa* | cefepime | 18 | 83.3% | 1.00 |
| *Pseudomonas aeruginosa* | ceftazidime | 21 | 52.4% | 0.88 |
| *Pseudomonas aeruginosa* | meropenem | 20 | 95.0% | 0.98 |

**Figure 4.**
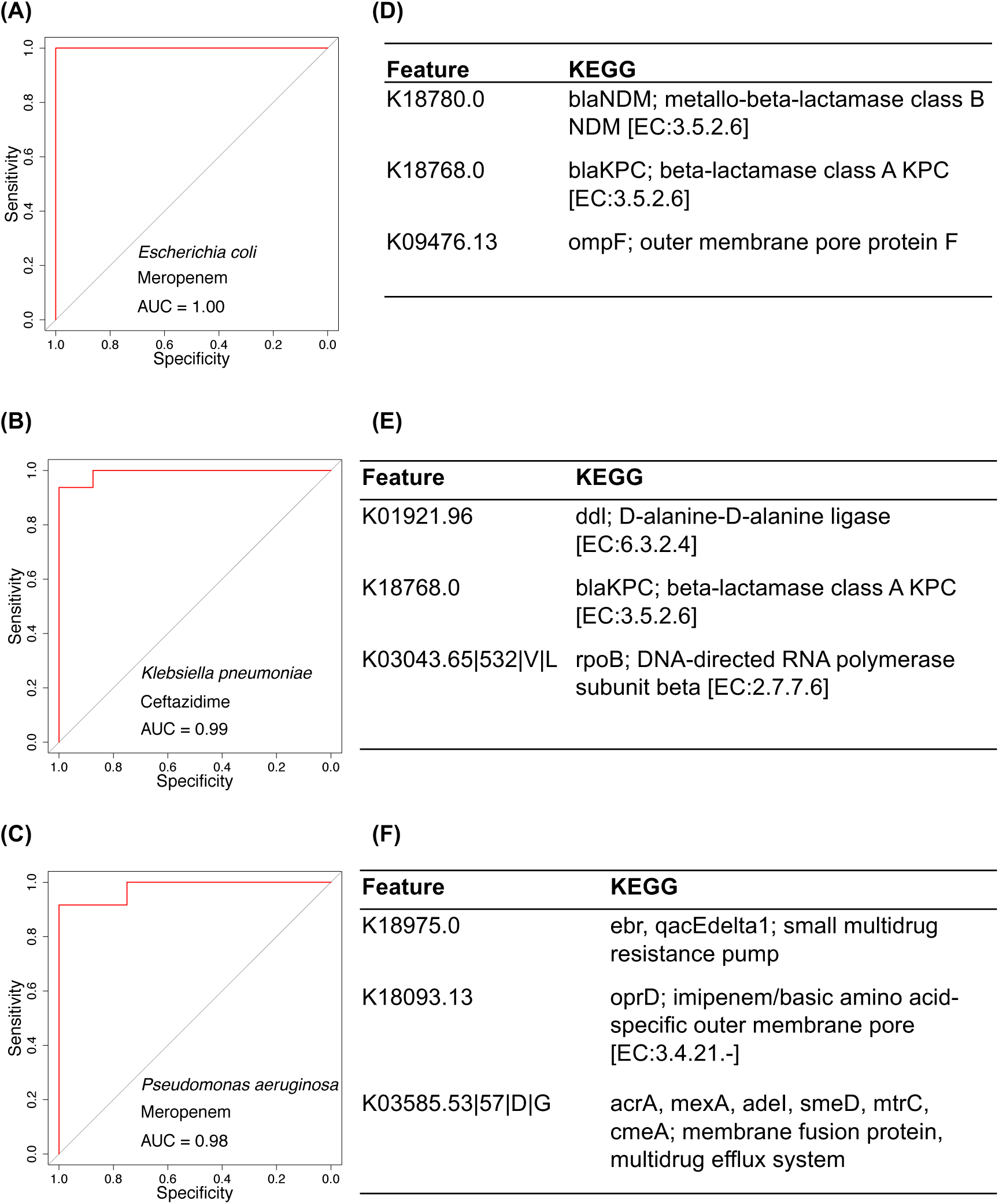
Validation performance metrics using an external dataset. AUROC (Area under the Receiver Operating Characteristic) for the prediction of the external dataset and top three predictors (KEGG orthlog variants based on importance) from the prediction models are reported.

**Figure 5.**
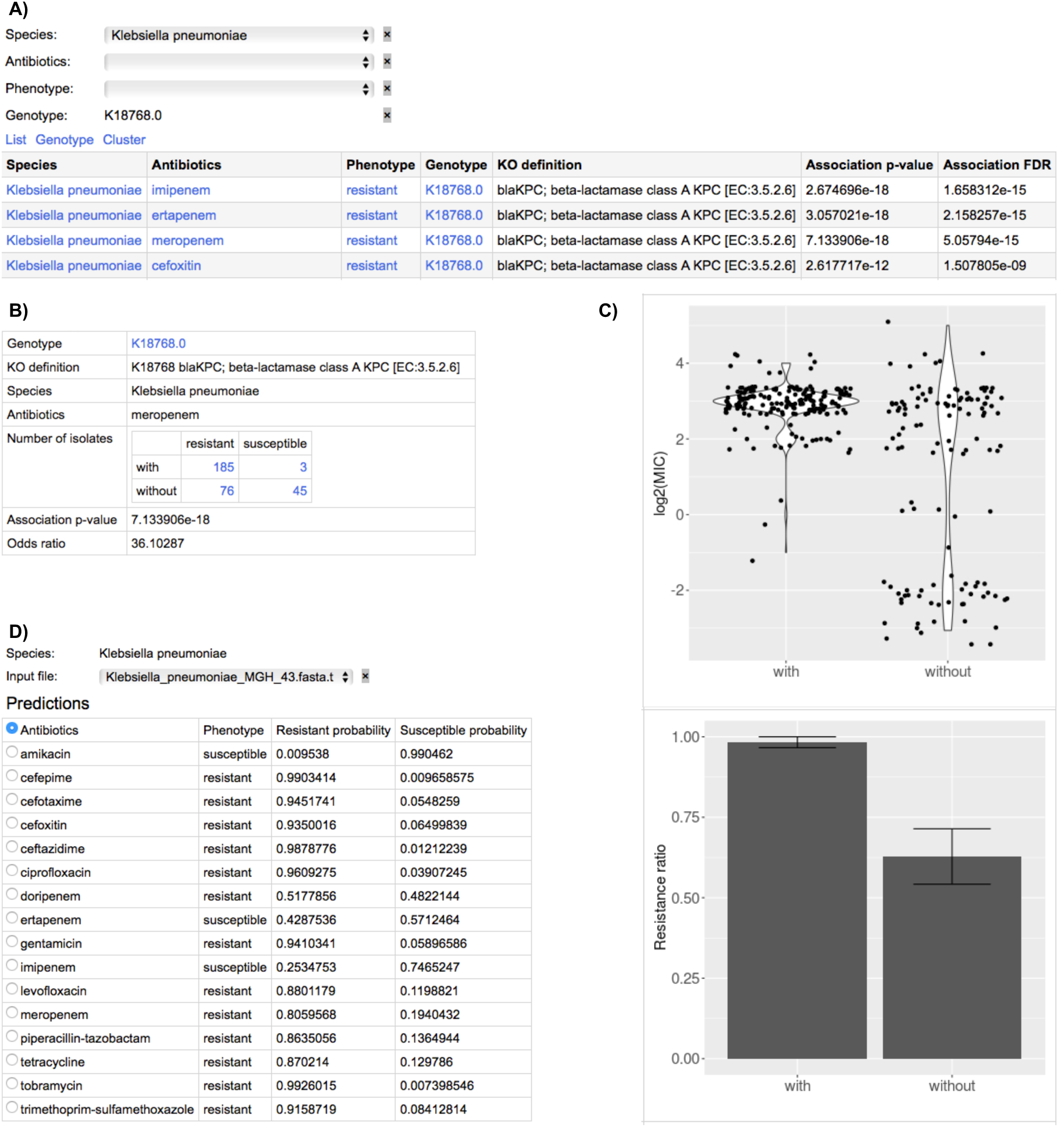
VAMPr provides rich sets of online resources for association models and prediction models. Users have the flexibility to explore known or novel antibiotic resistance-associated variants, and can upload their own sequence assembly and obtain predictions on antibiotic resistance. (A) association results webpage: users can explore variants, their interpretations, their statistical significance assessments; (B) detailed information, contingency table and odd-ratio for variant K18768 in the association model, and distribution plots; (C) Distribution plots for variant K18768 in the association model page; (D) prediction models allow for uploads of users’ sequence data for antibiotic resistance prediction.

### Offline resources for VAMPr pipeline

#### Offline resources – VAMPr source codes

We provide the source code that was used to create the association and prediction models. This allow users to curate and analyse their own sequence data for convenient offline usages. For example, users can provide FASTA sequence files and predict antibiotic resistance for multiple antibiotics without an internet connections.

## DISCUSSION

With the growing threat of antibiotic resistance and the rapidly decreasing costs associated with bacterial whole-genome sequencing, there is an opportunity for developing improved methods to detect resistance genes from genomic data (30). However, prior to the routine use of genomic data to routinely identify bacterial AMR status, there are several hurdles to be overcome including improving understanding of the genetic mechanisms underlying AMR for a broad-array of pathogen-antimicrobial combinations. To this end, we have developed the VAMPr pipeline to discover variant-level genetic features from NGS reads which then be correlated with phenotypic AMR data. We anticipate that with the continued generation of WGS data for numerous medically important pathogens, the widespread employment of VAMPr will assist with both clarifying associations between genomic data and AMR as well as developing new lines of AMR mechanism research.

An important advance of our study was our utilization of a novel approach to classifying variants based on gene orthologs. Our approach is different than other prediction models such as in PATRIC(8,31-33) which utilized the adaptive boosting (adaboost) algorithm. Our results were comparable or better in performance depending on the antibiotic-pathogen combination. In addition, this approach is in contrast to other popular ways for looking at gene variants such as k-mers (34). In the k-mer method, the frequency of k consecutive nucleotide or amino acid bases are counted as sequence features. Although the k-mer approach is straightforward to compute, it is hard to explain the k-mer in the context of genes, which requires extra analysis steps to interpret. To avoid these limitations, we instead utilized gene orthologs. By aligning the bacteria genomes with a group of consensus orthologous gene sequences, we can determine variants that are present for any particular AMR gene in a particular isolate. As the sequence variants are linked to ortholog genes, this approach can not only identify the presence or absence of known resistance genes, but can also give added insight into the impact of amino acid variants on various resistance phenotypes.

To understand how genetic variants were linked to AMR phenotypes, we built data-driven association models. We utilized a large collection of isolate sequence data from NCBI SRA and matching antibiotic resistance phenotypes reported in the NCBI BioSample Antibiogram. This allowed for an unbiased screening for statistically significant associations between genetic variants and specific antibiotics for a variety of pathogens. Thus, another strength of this study was the large data universe that these models were built upon with over 38,248 pathogen-antibiotic comparisons performed. Other groups have developed some similar tools, including recent efforts to predict AMR for drugs used in the treatment of *Mycobacterium tuberculosis*(35). An advantage of VAMPr over existing tools is its ability to analyse data from any bacterial species, providing that there are sufficient numbers of bacterial genomes-AMR phenotypic data to develop robust models. The publicly available nature of VAMPr and the NCBI Antibiogram means that the predictive models of VAMPr should significantly improve moving forward.

Our attempt to develop prediction models utilizing machine learning algorithms allowed for the identification of genes that are associated with resistance to a particular antibiotic in an unbiased fashion. This could allow for both confirmation of known resistance markers as well as a discovery tool to find novel genes that contribute to resistance. It is important to note that the genes and variants that we identified as predictive of resistance does not imply causation. These are correlations, and further work will be needed to see whether identified genes that are not currently known to contribute to resistance are biologically active or just mere bystanders with other causal genes. Future efforts will include testing whether some of these predicted genes or variants in genes are in fact biologically relevant.

There were other limitations of our study. Our attempt to validate the prediction models with a small number of isolates that were not included in the original training set illustrates particular challenges. There was clearly strain diversity in the recently sequenced isolates that was not fully represented in the available NCBI training set which impacted our ability to fully validate our prediction models. This indicates that there continues to be a need for increased genome sequencing that is more broadly representative certain pathogens. In addition, some pathogens such as *Pseduomonas aeruginosa* have a smaller number of genomes available in the NCBI dataset with paired antibiogram data available while other pathogens (such as *Salmonella)* have a large number of genomes with AMR phenotypes available. It is likely that increasing the number of genomes available for training purposes in pathogens like *Pseudomonas* will likely further improve our accuracy of the prediction model approach. For example, the recent study of *M. tuberculosis* resistance collected 10,290 samples and the large scale enabled accurate prediction of point mutations and antibiotic resistance(35). Our future efforts are aimed at further refining the VAMPr models to include larger numbers of isolates with a mixture of antibiotic susceptibility phenotypes. In conclusion, we are providing the VAMPr online resources for researchers to utilize in their efforts to better study and predict antibiotic resistance from bacterial whole genome sequence data. Widespread employment of VAMPr may assist with moving whole genome sequencing of bacterial pathogens out of the research lab setting and into the realm of clinical practice.

## AVAILABILITY

VAMPr is an open-source program and is available in the GitHub repository (https://github.com/jiwoongbio/VAMPr).

## ACCESSION NUMBERS

Not available.

## SUPPLEMENTARY DATA

Supplementary Data are available at NAR online.

## ACKNOWLEDGEMENT

We would like to acknowledgement Jessie Norris’s suggestions to improve this manuscript.

## TABLE AND FIGURES LEGENDS

Table 1. Prediction metrics for 32 VAMPr prediction models.

Table 2. External validation of VAMPr prediction model.

Figure 1. Overview of the VAMPr workflow.

Figure 2. Summary of significant variant associations and prediction accuracies from 93 species-antibiotics combinations.

Figure 3. Examples of variant-phenotype relationships determined by the association models.

Figure 4. Validation performance metrics using an external dataset.

Figure 5. VAMPr provides rich sets of online resources for association models and prediction models.

